# Isolation, ‘omics characterization and organotypic culture of alveolar type II pulmonary epithelial cells

**DOI:** 10.1101/2021.08.19.456947

**Authors:** Huaibiao Li, Moritz Schütte, Magdalena Bober, Torsten Kroll, Lucien Frappart, Ghina Bou About, Yu-Chieh Lin, Tania Sorg, Yann Herault, Christoph Wierling, Oliver Rinner, Bodo MH Lange, Aspasia Ploubidou

## Abstract

The alveolar type II (AT2) epithelial cell fraction includes the stem cells of the pulmonary alveoli, functioning in lung homeostasis and post-injury repair. AT2 cells have been characterized primarily *in situ*, in transgenic mouse models. We report a new methodology for their isolation, their “omics” characterization and stroma-cell-free organotypic culture. Our multi-omics analysis identified high expression of genes involved in oxidative phosphorylation and of AP-1 components, as well as new phosphorylation sites in AT2 biomarkers. Furthermore, we show that supplementation with KGF, FGF10 & HGF suffices for the *in vitro* proliferation of AT2 cells and formation of alveolar organoids, suggesting that AT2-based organotypic development depends on ligands of the c-Met and FGFR2 receptors. The reported methodology and in-depth molecular characterization provide new tools for the *in vitro* and *in vivo* functional analysis of pulmonary cells and of mouse models of lung disease.

## INTRODUCTION

Pulmonary alveoli are gas-exchange sac-like structures, lined by two epithelial cell types with distinct functions: Squamous Alveolar Type I (AT1) cells cover most of the surface area of the alveolar sacs, are apposed to blood capillaries and mediate gas exchange; whilst cuboidal AT2 cells secrete pulmonary surfactant proteins with biophysical and immunological functions in lung physiology (Fehrenbach, 2001; Guillot et al. 2013; Hogan et al. 2014).

Postnatally, the AT2 cell population includes stem cells, able to self-renew and differentiate into AT1 cells during lung homeostasis or in response to injury (Barkauskas et al. 2013; Desai et al. 2014; Zacharias et al. 2018). Disruption of the balance between self-renewal and differentiation of AT2 cells contributes to oncogenesis (Desai et al. 2014) and AT2 cells are thought to be at the cellular origin of lung adenocarcinoma (the most common type of Non-Small Cell Lung Cancer - NSCLC) (Mainardi et al. 2014; Sutherland et al. 2014). AT2 cell dysfunction can also lead to respiratory distress syndrome, chronic obstructive pulmonary disease or pulmonary fibrosis (Fehrenbach 2001).

The importance of AT2 cells in lung physiology has raised interest in their cell-intrinsic and microenvironment-induced regulation, in particular under pathological conditions - including upon infection with SARS-CoV-2. Accordingly, omics characterization of AT2 cells has mostly addressed post-infection analysis (Seddigh et al. 2017; Shiraishi et al. 2019a; Shiraishi et al. 2019b; Zacharias et al. 2018).

Mouse *in vivo* studies employing transgenic and lung injury models as well as organotypic systems of the pulmonary epithelium and, more recently, iPSC-approaches, have enabled alveolar cell lineage and differentiation analysis (reviewed in Barkauskas et al. 2017; Beers and Moodley, 2017; Hogan et al. 2014). AT2 cells form alveolar organoids when co-cultured with fibroblasts in Matrigel-based extracellular matrix (ECM), reflecting the essential role of stromal cells *in vivo,* in modulating AT2 functions (Barkauskas et al. 2013). The growth factors, secreted by stromal cells to support *in vitro* AT2 cell growth, differentiation and alveolar organoid formation, remain to be identified. In this regard, a simplified organotypic culture method, without stromal cells, would be advantageous in dissecting the distinct role of growth factors in AT2 self-renewal and differentiation.

We report (i) a modified method of AT2 cell isolation; (ii) the quantitative transcriptomic and proteomic characterization of primary AT2 cells, including new phosphorylation data of core AT2 cell components and of proteins relevant to SARS-CoV-2 infection; (iii) the identification of signaling pathways that promote the proliferation of AT2 cells and the formation of alveolar organoids in stromal cell-free culture.

## RESULTS & DISCUSSION

Isolation of primary cells for omics analysis (and *in vitro* differentiation) relies on bulk purification methodology to yield adequate amount of enriched material, albeit one that minimizes isolation-introduced artefacts, which would elicit changes in gene expression or protein activation state of the target cells (He et al. 2018). This is particularly important when the expression profiling investigates dynamic states of signaling pathways.

Primary AT2 cell isolation for organotypic culture (Table S1) has been based on affinity purification via AT2-specific cell surface markers followed by fluorescence-activated single cell sorting (FACS) and yields target cells of high purity. Nonetheless, the FACS-associated mechanical, hydrodynamic and electrical stress can adversely affect several physiological cell functions, including viability, of the sorted cells (Holt and Olsen, 2016; Li et al. 2013). Compared to FACS, magnet-assisted cell separation (MACS) (Miltenyi et al. 1990) is gentler (Bowles et al. 2019), thereby resulting in higher cell viability (Sutermaster and Darling, 2019).

To obtain an enriched fraction of mouse primary AT2 cells for comprehensive omics characterization and organotypic culture, we devised an antibody- and magnetic sorting-based methodology, building on previously published protocols (Table S1) (Barkauskas et al. 2017).

### AT2 cell isolation

Antibody-based affinity purification of rat and human AT2 cells can utilize surface markers recognized by antibodies (RT2-70, HT2-280) of unknown molecular specificity that do not, however, cross-react with mouse antigens (Gonzalez et al. 2005; Gonzalez et al. 2010). To overcome this obstacle in murine AT2 cell isolation, a fluorescent reporter gene driven by an AT2-specific promoter (Sftpc-CreER) (Barkauskas et al. 2013), or EpCAM (a transmembrane epithelial surface protein) (Hasegawa et al. 2017) have been used as marker of AT2 mouse cells (Table S1). The former method (genetic introduction of reporter and promoter constructs in the mouse lines of interest) is time-, labor- and cost-intensive. A reporter- and promoter-independent method for AT2 cell isolation would, thus, be preferable.

To circumvent these issues, we used EpCAM, the only known surface protein of murine AT2 cells, in their positive selection (Fig.S1). To prevent cross-contamination with AT1 cells, that also express EpCAM (Hasegawa et al. 2017; Kasper et al. 1995), we first depleted the lung cell population of AT1 cells, in a negative selection step, using an antibody against their surface marker T1α (Williams, 2003) (Fig.S1). An anti-CD45 antibody was included in the negative selection to deplete immune cells which are abundant in the lung (Happle et al. 2018).

Following negative selection, both T1α/CD45-positive (+ve) and T1α/CD45-negative (−ve) cell populations were analysed by immunofluorescence microscopy using the AT2 cell specific marker proSPC (Surfactant Protein C) (Fehrenbach, 2001). The majority of proSPC-expressing cells remained in the T1α/CD45-ve fraction (Fig.1A), which was subsequently subjected to positive selection with an anti-EpCAM antibody (Fig.S1). The yield attained (2-3×10^6 EpCAM expressing cells/mouse lung) is comparable to that obtained by a FACS-based similar protocol (Sinha and Lowell, 2016b).

**Figure 1:**
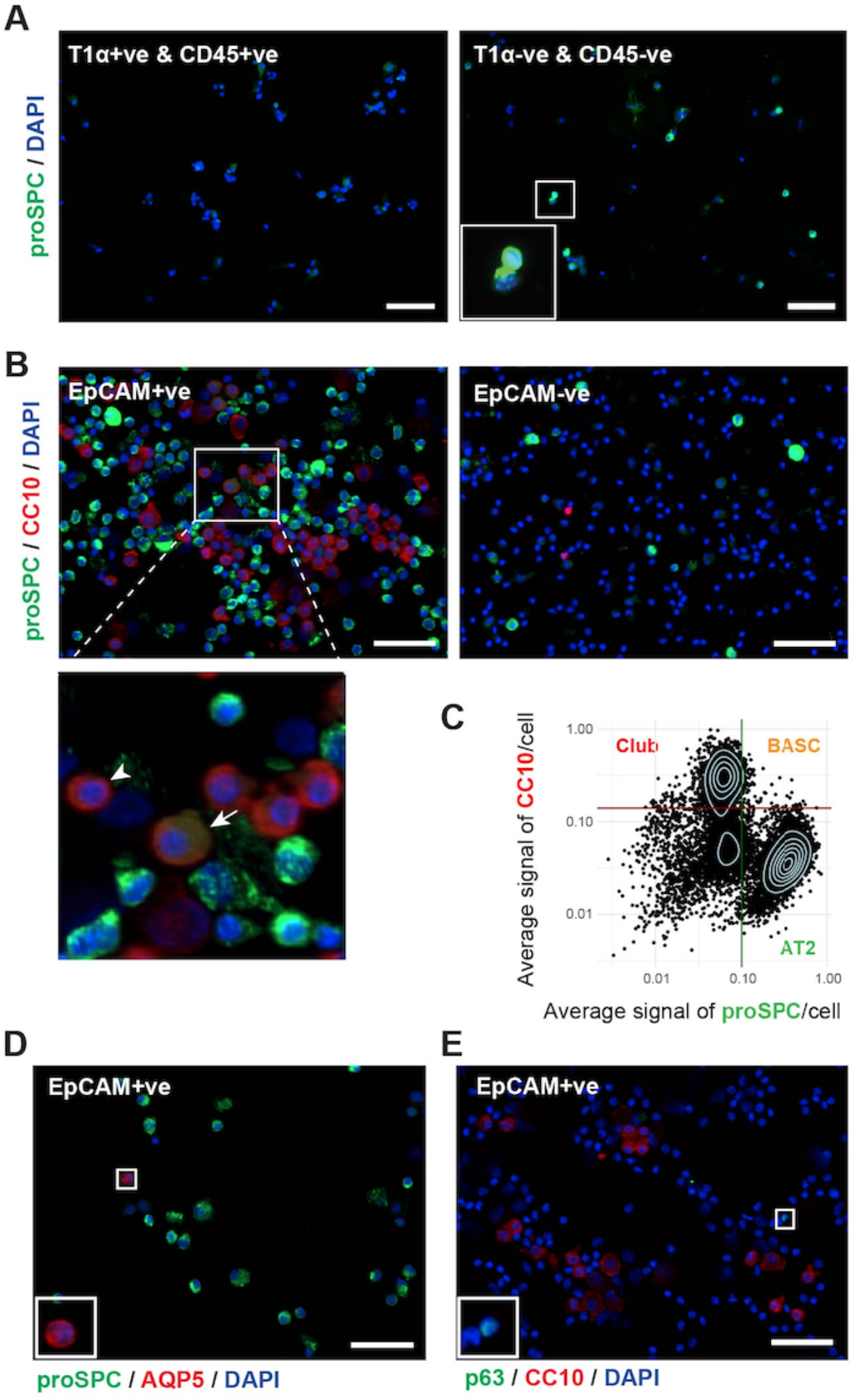
Enrichment of AT2 cells. Immunofluorescence microscopy micrographs depict the cells of the indicated column fractions, labeled with the indicated antibodies and DAPI, to identify and quantify different cell types. (**A, B, E, F**). Example of dot plot of average intensity fluorescence per cell (proSPC *versus* CC10, in the EpCAM+ve fraction) (**C**). Thresholds set for proSPC (green line) and CC10 (red line) are indicated. AT2, BASC and club cell populations are indicated. In addition to AT2 cells, the (EpCAM+ve) column eluate contained club cells (**B arrowhead**) and BASCs (**B arrow**), plus small numbers of AT1 (**D**) and basal cells (**E**) – all shown in the magnified inserts. Scale bars: 50μm.

Following positive selection, we analyzed the identity and relative abundance of different cell types, within the EpCAM+ve fraction, via immunofluorescence co-labelling and High-Content Analysis (HCA) microscopy, using antibody markers for AT2 (anti-proSPC), club (anti-Scgb1a1, also known as anti-CC10), basal (anti-p63) and AT1 (anti-AQP5) cells (Fig.1, *Suppl. Experimental Procedures*). For HCA-based quantification of cell subtypes, the thresholds were set based on the distribution of average intensities, as illustrated by the dot plot (Fig.1C) for proSPC and CC10 double-labelled cells.

In addition to the alveolar proSPC+ve cell population (Fig.1B,C; Table S5), the EpCAM+ve fraction also contained proSPC-ve cells. The latter are likely bronchiolar epithelial cells – as both basal cell progenitors (Tata et al. 2013) and club cells (McQualter et al. 2010) of the bronchioles express EpCAM. Indeed, most proSPC-ve cells were CC10+ve (Fig.1B arrowhead; Table S5), in agreement with a previous study (Shiraishi *et al.* 2019a). Notably, ca. 1.8% of proSPC+ve cells also express CC10 (Fig.1B arrow, Fig.1C, Table S5), a characteristic of Bronchio-Alveolar Stem Cells (BASC) (Kim et al. 2005; Liu et al. 2019; Salwig et al. 2019). Finally, a small number of AT1 (Fig.1D) and basal cells (Fig.1E; Table S5) were co-isolated.

We conclude that, until murine AT2 cell-specific surface markers are identified, a succession of T1α (negative), CD45 (negative) and EpCAM (positive) selections applied via MACS can be used to isolate AT2 cells. This method inadvertently co-isolates BASCs and bronchiolar epithelial cells along with the AT2 cell population. For isolation of rat or human AT2 cells using this protocol, replacement of the EpCAM antibody with RT2-70 or HT2-280, respectively (Gonzalez et al. 2005; Gonzalez et al. 2010), would prevent contamination with bronchiolar cells, thereby increasing enrichment of the AT2 cell fraction.

### Transcriptomic, proteomic & phospho-proteomic characterization

The EpCAM+ve lung cells, isolated from two cohorts of wild type mice [either untreated (WT) or intranasally injected with an adenoviral vector encoding the Cre recombinase under the control of the Sftpc promoter (WT^Ad-SPC-Cre^)], were further characterized by transcriptome and proteome analysis.

Overall, 13.274 transcripts were identified and quantified by RNA sequencing, and 5.952 proteins by LC-MS (Tables S2 & S3) – a significantly larger dataset than previously reported for these cells (Treutlein et al. 2014). Among the translated transcripts (Fig.2A,B), the most highly expressed were surfactants (Sftpa, Sftpb, Sftpc) and Lyz2 - all known AT2 cell markers (Fehrenbach, 2001) - thus confirming AT2 cell enrichment in both WT (Fig.2A) and WT^Ad-SPC-Cre^ (Fig.2B) derived EpCAM+ve lung cells. Expression of Scgb1a1 concurs with the HCA data (Fig.1C) identifying a sub-population of double proSPC+ve and CC10+ve cells, within the EpCAM+ve cells.

**Figure 2:**
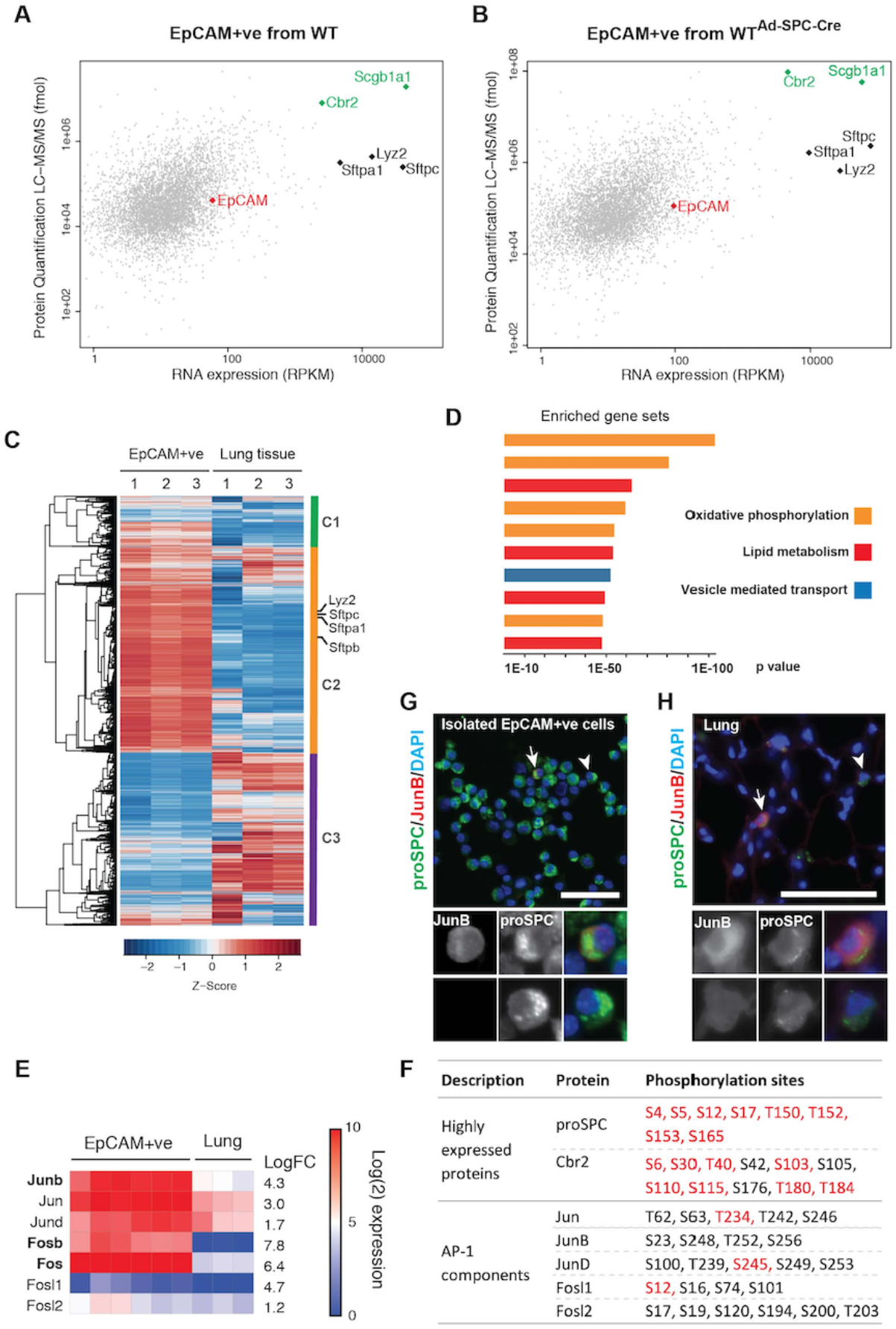
‘Omics characterisation of AT2 cells. (**A**, **B**) Dot plots of mRNA versus protein expression of lung EpCAM+ve cells isolated from the indicated mouse lines. Highly expressed genes (AT2 cell markers in black, BASC/club cell markers in green) and EpCAM are indicted. (**C**) Gene expression heatmap comparing RNA profiles of EpCAM+ve cells *versus* adult lung tissue samples for the genes expressed also at protein level; values are scaled per row. Three gene clusters, differentially expressed between isolated cells and lung tissue, are indicated on the right. AT2 cell markers, indicated, are highly over-represented in the isolated cell fraction (cluster C2). (**D**) GSEA for genes of cluster C2 has identified, among the top 10 enriched gene sets, pathways operating in oxidative phosphorylation, lipid metabolism and vesicle-mediated transport. NES: Normalized Enrichment Score. (**E**) Gene expression heatmap and fold change of AP-1 transcription factor subunits in isolated cells *versus* lung tissue. (**F**) Phosphorylation sites of highly-expressed proteins and of AP-1 subunits, detected by MS analysis. Newly identified (red) and known (black) phosphorylated residues are indicated. Immunofluorescence labelling of JunB in AT2 cells, isolated (**G**) or *in situ* in the lung (**H**). Arrows indicate JunB+ve and arrowheads JunB-ve AT2 cells. Scale bars: 50μm (**G,H**).

Among the most abundant proteins, several new post-translational modifications in Cbr2 and proSPC were identified by mass spectrometry (Fig.2F, Table S3). Sftpc encodes the Surfactant Pulmonary protein C (SPC) which is primarily secreted by AT2 cells (Beers & Moodley 2017). Secreted pulmonary surfactant proteins have important biophysical and immunological functions in lung physiology. This surfactant is synthesized as the precursor proSPC by AT2 cells and subsequently proteolytically cleaved to SPC; the latter is secreted via the lamellar bodies (Keller et al. 1991). Intracellular processing of proSPC requires the region spanning residue Met10 to Thr18; its deletion results in retention of proSPC in the ER (Johnson et al. 2001). Within this propeptide region of proSPC, we identified phosphorylation of Ser12 and Ser17 (Fig.2F). The BRICHOS domain at the C-terminus of proSPC is critical for polypeptide folding during the intracellular processing of the precursor. Mutations at this domain can trigger the formation of proSPC cellular aggregates, thereby activating the unfolded protein response (Mulugeta et al. 2007). Within the BRICHOS domain, we identified phosphorylation in Thr150, Thr152, Ser153 and Ser165 (Fig.2F). Given the established link between Sftpc dysfunction and lung fibrosis (Brasch et al. 2004; Nureki et al. 2018), the impact of these phosphorylation events on the subcellular localization and secretion of SPC warrants further investigation.

We subsequently performed differential gene expression analysis to identify pathways critical for AT2 function. To this end, we sequenced the whole lung tissue of WT^Ad-SPC-Cre^ mice, and compared the expression profile of lung tissue to that of the EpCAM+ve cell fraction.

First, clustering analysis of genes expressed at both transcript and protein level, detected by RNA-Seq and MS methodologies, revealed striking differences in expression pattern between the lung and isolated cells from WT^Ad-SPC-Cre^ mice (Table S2, Fig.2C). Established AT2 cell markers (e.g. Sftpc, Lyz2) are found in the cluster (C2) of genes that are highly enriched in the EpCAM+ve cell population, when compared to the lung (Fig.2C). Second, we sought to identify cell-surface proteins that could be used in direct isolation of mouse AT2 cells. Using LogCPM>1 and FDR<0.05 as threshold, from 109 transcripts highly enriched in the EpCAM+ve fraction seven (Table S3) are predicted to encode cell surface proteins (Cell Surface Protein Atlas database, Omasits et al. 2014). Nevertheless, the corresponding proteins were not detected in the proteome, suggestive of low-level translation.

We analyzed the signaling pathways enriched in cluster 2 (C2) (Fig.2C) via Gene Set Enrichment Analysis (GSEA). The top 10 enriched gene sets include components of fatty acid metabolism and vesicle-mediated transport (Fig.2D), in agreement with previous reports that AT2 cells are the major source of surfactant proteins within the alveolar region (Shiraishi et al. 2019b; Treutlein et al. 2014). Additionally, components of pathways associated with the function of mitochondria, such as oxidative phosphorylation and respiratory electron transport, are highly enriched in cluster 2 (Fig.2D). Notably, these metabolic pathways are significantly deregulated in iPSC–derived AT2 cells (iAT2) upon SARS-CoV-2 infection (Hekman et al. 2020).

Altogether, the GSEA results highlight the importance of mitochondria in AT2 cell homeostasis. As mitochondria generate reactive oxygen species (ROS), we examined the expression of ROS-related proteins. Among components of the ROS pathway, JunB – a subunit of the AP-1 transcription factor – is highly enriched in the AT2 cell fraction (Fig.2E). Indeed, single cell RNAseq analysis has identified Jun as key transcription factor in the metabolic pathway of mature AT2 cells (Treutlein et al. 2014). JunB forms heterodimers with other AP-1 family members (Eferl and Wagner, 2003). Among these, c-Fos and Fosb transcripts were also markedly enriched in AT2 cells (Fig.2E). Furthermore, phospho-proteome analysis detected multiple phosphorylation sites in AP-1 family proteins (Fig.2F, Table S3). The activity and turnover of AP-1 components is tightly regulated by phosphorylation (Boyle et al. 1991; Papavassiliou et al. 1992). For instance, phosphorylation at S251, T255 and S259 triggers degradation of human JunB by FBXW7 (Perez-Benavente et al. 2013). Our analysis showed that the corresponding conserved domain of mouse JunB, phosphorylation of which targets the protein for degradation (phospho-degron), is phosphorylated (S248, T252, S256) in the EpCAM+ve cell fraction (Fig.2F). Moreover, both in isolated AT2 cells as well as *in situ* in the lung, JunB is localized in the cytoplasm, not in the nucleus (Fig.2G,H). These results collectively suggest that phosphorylation at the phospho-degron sites of JunB promotes its degradation in the cytoplasm, at homeostatic state.

Recent investigations in SARS-CoV-2, using both established and new infection systems, have mapped the differential phosphorylation of host proteins during infection. In iAT2 cells, neither phosphorylation of proSPC nor of JunB are affected by SARS-CoV-2; nonetheless, only one proSPC phospho-site (Ser5) was detected and quantified (Hekman et al. 2020). Here, using MACS-based enrichment of AT2 cells and quantitative mass spectrometry, we have identified several new phosphorylation sites in proSPC and in members of the Jun family (Fig.2F), as well as in several SARS-CoV-2-interacting proteins (Table S3) that were previously identified in kidney cells (Bouhaddou et al. 2020). We suggest that the new phospho-sites identified here represent the undisturbed state of intracellular signaling in AT2 cells; it remains to be investigated whether and how they are modulated by SARS-CoV-2.

### Organotypic culture

To functionally test the proliferation and differentiation capacity of the EpCAM+ve lung cells, we cultured them in the ECM-equivalent Matrigel, under several different conditions (Fig.S1B). AT2 cells can form spheroids when co-cultured with lung fibroblasts (Barkauskas et al. 2013; Jain et al. 2015; Ng-Blichfeldt et al. 2018). Alternatively, the simultaneous *in vitro* modulation of multiple signaling pathways (including Wnt, TGF-beta, BMP, Notch, FGFR2) enables formation of alveolospheres without the support of fibroblasts (Shiraishi et al. 2019a; Shiraishi et al. 2019b; Youk et al. 2020; Katsura et al. 2020). To identify, among the above candidates, the pathway/s responsible for (i) AT2 proliferation and (ii) alveolosphere formation, we systematically tested the effect of their modulation in fibroblast-free organotypic culture.

#### 1. Tracheospheres & bronchioalveolar organoids

The pool of AT2 stem cells is maintained by Wnt5a, which is secreted by the fibroblast niche (Nabhan et al. 2018). Two-week culture of the EpCAM+ve lung cells in Matrigel overlaid by MTEC/plus media containing Wnt5a ligand (Fig.3A) gave rise to spheroids, additionally forming pseudostratified epithelium (Fig.3B) reminiscent of tracheospheres derived from basal cells (Rock et al. 2009). Expression of p63 in cells at the basal layer of the organoids (Fig.3C), supports the conclusion that basal cells had amplified and gave rise to **tracheospheres** under these culture conditions (Barkauskas et al. 2017; Rock et al. 2009; Sachs et al. 2019). Few proSPC+ve cells were observed in these cultures, mostly as single cells (Fig.3D, arrows), suggesting that Wnt5a alone does not suffice to drive proliferation of AT2 cells.

**Figure 3:**
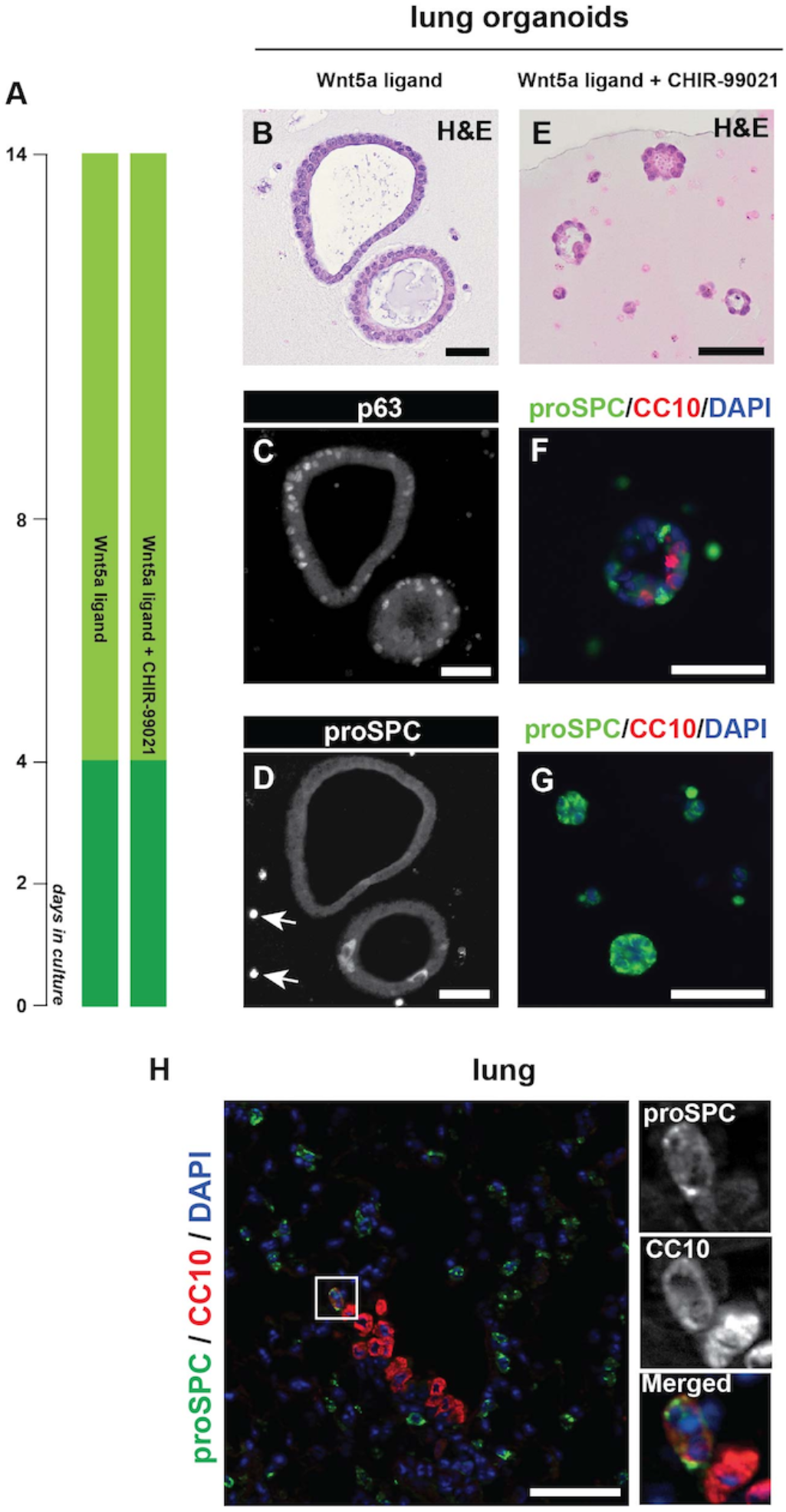
Activation of Wnt pathway induces the formation of tracheospheres and bronchioalveolar organoids. (**A**) Schematic representation of culture conditions (see also Fig.S1). Haematoxylin and Eosin (H&E) stained (**B, E**) or immunofluorescence-labelled (**C, D, F,G**) sections of organoids cultured in the presence of Wnt5a (**B, C, D**), or Wnt5a and CHIR-99021 (**E, F, G**). Wnt5a supplementation induced formation of tracheospheres comprising p63+ve basal cells (**C**), while AT2 cells (arrows) did not proliferate (**D**). Wnt5a and CHIR-99021 together induced development of bronchioalveolar organoids composed by club and AT2 cells (**F**) or alveolar organoids of AT2 cells (**G**). (**H**) Immunofluorescence labelling of BASCs in the lung, co-expressing AT2 and club cell markers, residing at the junction area between bronchioles and alveoli. Scale bars: 50μm.

This result was rather surprising, given that lung fibroblast-secreted Wnt5a is reported to maintain the stemness of AT2 cells (Nabhan et al. 2018). We therefore included in the culture media CHIR-99021, a small molecule inhibitor of GSK3 (Ring et al. 2003), together with the Wnt5a ligand, to enhance activation of the Wnt pathway. Under these conditions, small lumen-forming spheroids developed, comprising a single layer of cuboidal cells (Fig.3E) that express either proSPC or the club cell marker CC10 or both proteins (Fig.3F). The observed co-expression of AT2 and club cell markers is only encountered in Broncho-Alveolar Stem Cells (BASCs) located at the Broncho-Alveolar Duct Junction (BADJ) (Fig.3H). Given the increasing evidence that BASCs can differentiate into either pro-SPC+ve or CC10+ve cells (Kim et al. 2005; Liu et al. 2019; Salwig et al. 2019), we infer that these spheroids are **bronchioalveolar organoids** (McQualter et al. 2010; Ng-Blichfeldt et al. 2018) derived from BASCs present in the EpCAM+ve lung cell fraction (Fig.1B,C).

In addition to the bronchioalveolar organoids, small cell clusters with barely distinct lumen were formed under these culture conditions, comprising exclusively proSPC+ve AT2 cells (Fig.3G). We conclude that activation of the Wnt pathway triggers expansion of AT2 cells. The small size (<40μm in diameter) and morphology of these AT2 cell clusters suggest that they are at an early stage of organotypic growth.

#### 2. Alveolospheres

Rapid expansion of AT2 cells is especially important during alveolar regeneration after lung injury. FGFR2 ligands (e.g. KGF and FGF10) are necessary for the proliferation of AT2 cells during lung repair (Ulich et al. 1994; Yano et al. 2000; Yuan et al. 2019). What are the *in vitro* requirements?

In co-culture with fibroblasts, supplementation with FGF10 and HGF enhances colony formation of EpCAM+ve cells (McQualter et al. 2010). In the absence of support cells, KGF is necessary but not sufficient for AT2 spheroid formation (Shiraishi et al. 2019a); additional culture supplements are thus required (Table S1).

We therefore tested whether the FGFR2 ligands **K**GF, **F**GF10 and **H**GF (hereafter called **KFH**) suffice to induce amplification and spheroid culture of AT2 cells, in the absence of stromal cells (Fig.4A). In the absence of KFH (Fig.4B), the inoculated single AT2 cells remained proSPC+ve but they did not amplify (PCNA-ve), as anticipated (Barkauskas et al. 2013). In contrast, addition of KFH induced AT2 cell expansion (proSPC+ve and PCNA+ve) and their organization into spheroids (Fig.4C). These results suggest that AT2 cell proliferation is positively regulated by FGFR2 and the c-MET pathway.

**Figure 4:**
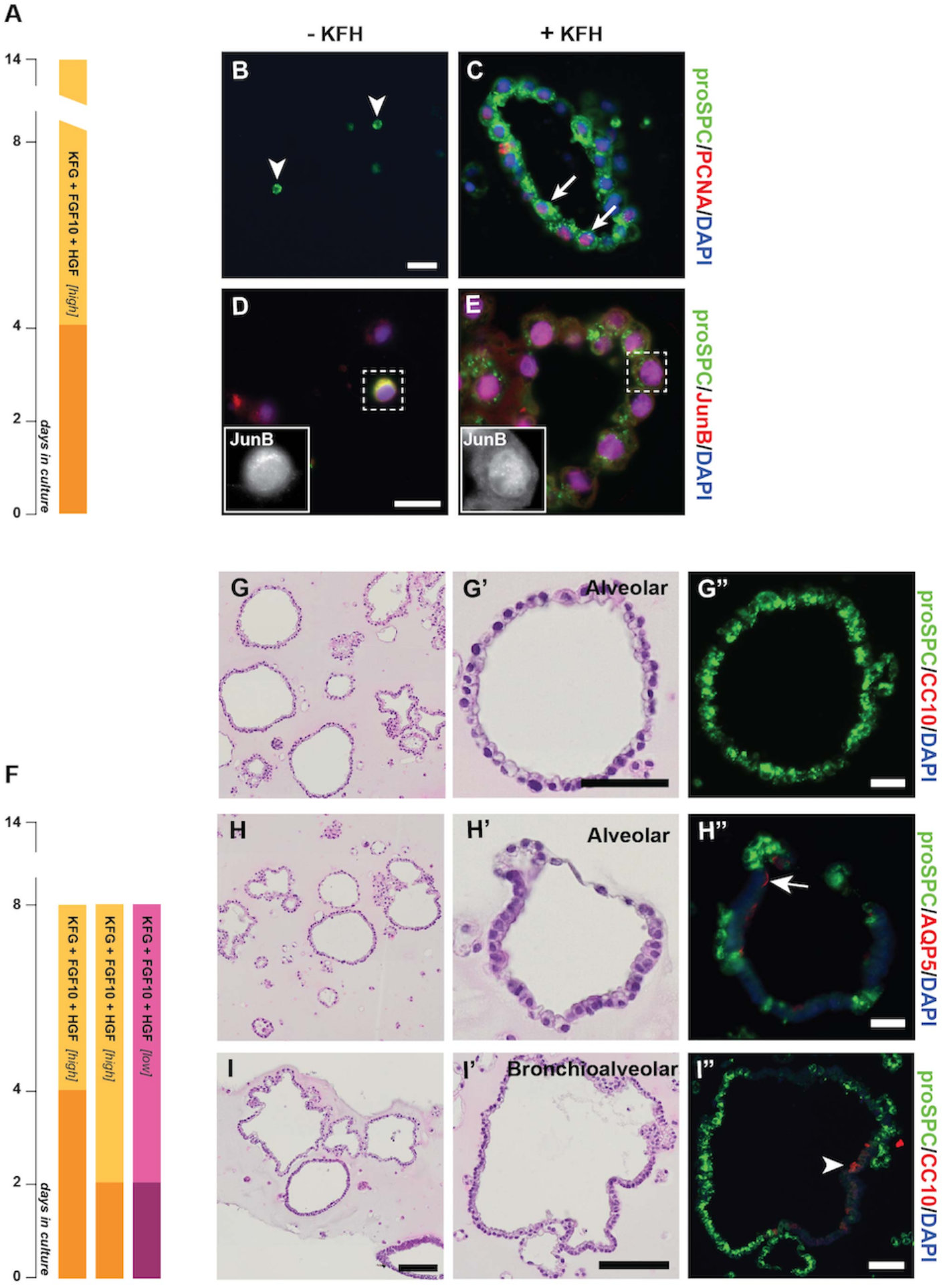
KGF, FGF10 and HGF synergistically promote the formation of bronchioalveolar and alveolar organoids. (**A, F**) Schematic representation of culture conditions (see also Fig.S1). Immunofluorescence labelling of sections from cultures grown with (**C, E**) or without (**B, D**) KFH. In the latter condition, AT2 cells did not proliferate (**B**, arrowheads) and exhibited cytoplasmic localization of JunB (**D**) indicative of quiescence. Supplementation with KFH induced AT2 cell proliferation, demonstrated by expression of PCNA (**C**) and nuclear localization of JunB (**E**), and organoid formation. H&E stained (**G, G’, H, H’, I, I’**) and immunofluorescence labeled (**G”, H”, I’’**) semi-serial sections of 8-day organoids cultured as indicated in **F** (panels **G** and **H** from high KFH cultures; **I** from low KFH). Most organoids developed were alveolospheres, either comprising exclusively AT2 cells (**G’, G”**) or AT2 plus AT1 cells (**H’, H”**). Additionally, bronchioalveolar organoids (**I’, I”**; arrowhead indicates club cell) and tracheospheres also formed. Scale bars: 20μm (**B**, **C**, **D, E**), 50μm (**G**, **H**, **I**, **G’**, **H’**, **I’**, **G”**, **H”**, **I”**)

Further experiments demonstrated that eight-days culture of the EpCAM+ve cell fraction (Fig.4F), in the presence of KFH, was sufficient to induce cell expansion and their polarization into lumen-forming organoids with size of ca. 50-200μm in diameter comprising cuboidal cells (Fig.4G,H,I). We compared two concentrations of KFH (Fig.S1B, *Experimental Procedures*); the higher one led to increased number and diameter of organoids formed.

Light and immunofluorescence microscopy revealed four distinct organoid subtypes. The majority (ca. 80%) were **alveolospheres**:

i. Alveolospheres comprising exclusively proSPC+ve AT2 cells (Fig.4G’,G”). Their development suggests that KFH promotes AT2 cell amplification and polarization.
ii. Alveolospheres containing AT2 cell clusters flanked by AQP5+ve cells (an AT1 cell marker) (Fig.4H’,H”). As lineage tracing studies have demonstrated that AT2 cells differentiate to AT1 cells (Barkauskas et al. 2013), we postulate that this type of organoid also arises from AT2 cells. In addition, two other organoid types were formed:
iii. Bronchioalveolar organoids comprising proSPC+ve AT2 cells, CC10+ve club cells and cells expressing both markers (Fig.4I’,I”). These organoids differ greatly in size to the bronchio-alveolar ones generated under Wnt activation conditions (Fig.3E,F *cf*. Fig.4I’,I”).
iv. Tracheospheres likely derived from basal cells.

Extending the culture time from 8 to 14 days (Fig.S2 F) led to two outcomes: the cuboidal cell organoids (Fig.S2 B,C arrows) became largely proSPC-ve (Fig.S2 D,E), whilst large cell clumps (Fig.S2 B,C arrows) comprising vacuolated and partly proSPC+ve cells (Fig.S2 D,E) also formed.

We found that expression of JunB is enriched in the EpCAM+ve cells (Fig.2E). Does KFH-induced proliferation affect its activity? As component of the AP-1 transcription factor, active JunB localizes in the nucleus to promote cell cycle progression (Farras et al. 2008). However, in quiescent AT2 cells, JunB is cytoplasmic (Fig.2G,H). During organotypic culture of these cells, in the absence of KFH JunB remained in the cytoplasm (Fig.4D), translocating in the nucleus upon KFH addition in the media (Fig.4E).

We conclude that supplementation of mouse primary AT2 cells with FGFR2 ligands is necessary and sufficient for their proliferation and differentiation into alveolospheres, independently of fibroblasts.

#### 3. In vitro proliferation & differentiation of AT2 cells – Growth Factors

Our findings raised the question whether FGFR2 ligands regulate proliferation or differentiation of AT2 cells. The available evidence suggests direct regulation of the former. First, primary AT2 cells proliferate when grown atop a Matrigel-and-lung fibroblast matrix (Sucre et al. 2018). Second, lung fibroblast-conditioned medium (which contains KGF and HGF, the primary mitogens for AT2 cells) stimulates the proliferation of rat AT2 cells (Panos et al. 1993). Nonetheless, KGF is reported to prevent the differentiation of AT2 into AT1 cells (Qiao et al. 2008). Indeed, our cultures (initiated with an enriched AT2 cell population but negligible number of AT1 cells) gave rise to relatively few organoids comprising both AT2 and AT1 cells (Fig.4G’,G’’). We therefore could not exclude that continuous exposure to these growth factors blocks differentiation of AT2 cells to AT1.

To test this supposition, we removed KFH at day 8, and cultured the organoids for another 6 days (Fig.S2 F). These culture conditions did not increase the AQP5+ve cell population (Fig.S2 G–J), suggesting that removal of KFH is not sufficient to induce the differentiation of AT2 cells. We cannot rule out the possibility that KFH had diffused into the Matrigel and was not completely removed by the change of the overlaying culture media, nor that that the simultaneous removal of FGF10 and HGF, together with KGF, played a role.

Hence, in fibroblast-free culture conditions, FGFR ligands appear to promote the rapid proliferation of AT2 cells. Further modulation of the relative concentration of the 3 ligands, during organotypic culture, has the potential to increase the efficiency of *in vitro* differentiation of AT2 cells into AT1.

#### 4. In vitro proliferation & differentiation of AT2 cells – Niche

Within the alveoli, lung fibroblasts likely serve as the source of FGFR2 ligands. Hence, replacement of fibroblasts with KFH, *in vitro,* stimulates AT2 cell proliferation and formation of alveolar organoids. However, noteworthy are the morphological and structural differences between alveolospheres generated in the presence of mesenchymal cells (Barkauskas et al. 2013; Jain et al. 2015; Ng-Blichfeldt et al. 2018; Shiraishi et al. 2019b) and their counterparts cultured without supporting cells (Fig.4). In the former, an interior cluster of AT1 cells is surrounded by AT2 cells, while fibroblasts spread throughout the organoid (Barkauskas et al. 2017; Barkauskas et al. 2013). In the latter, the alveolospheres comprise a single layer of epithelial cells (Fig.4).

These differences highlight the importance of the stem cell niche in modulating AT2 functions. AT2 cells do not differentiate to AT1 cells when grown in close contact with lung fibroblasts (Sucre et al. 2018), suggesting that the proximity to mesenchymal cells is critical to maintain the stem cell state of AT2 cells. Factors secreted by mesenchymal cells, such as Wnt and FGFR2 ligands, are plausible gatekeepers of the stem cell state (Nabhan et al. 2018; Shiraishi et al. 2019a).

Notably, the diameter of organoids generated under KFH stimulation (Fig.4) is significantly larger than of those cultured under Wnt pathway activation (Fig.3) suggesting that the impact of the FGFR2 pathway on BASC and AT2 cell proliferation is much stronger than that of the Wnt pathway. The differential response of AT2 cells to different stimuli reflects the distinct role of these signals in maintaining alveolar homeostasis. Activation of the Wnt pathway has a mild effect on AT2 proliferation; its major function is to maintain the AT2 stem cell pool in the quiescent alveoli as well as during alveoli regeneration (Nabhan et al. 2018; Zacharias et al. 2018). Upon lung injury, however, lung mesenchymal cells increase the expression of FGFR ligands to trigger the bulk expansion of AT2 cells. Likely, the AT2 cells exposed to Wnt ligands from neighboring mesenchymal cells maintain their stem cell identity, while their counterparts away from niche lose the signal of FGFR2 ligands and differentiate into AT1 (Nabhan et al. 2018).

## Conclusions

Using multi-omic analysis downstream of gentle isolation methodology, we identified the gene sets with enriched expression in AT2 cells as well as new phosphorylation sites of AT2 biomarkers – altogether reflecting the unperturbed expression profile and signaling state of these cells. Furthermore, using stroma-free organotypic culture, our experiments pinpointed the key growth factors that drive alveolar organoid formation, hence facilitating the investigation of alveolar physiology *in vitro*. Indeed, it was recently shown that alveolar organoids infected with SARS-CoV-2 recapitulate the pathological changes observed in COVID-19-diseased lungs (Katsura et al. 2020) – thus demonstrating the suitability of this system in infection studies.

Inadvertently, the isolation strategy described here co-purifies AT2 cells and BASCs (Fig.1). What conclusions could be drawn for the *in vitro* differentiation of these cell types?

First, although club cells (CC10+ve) are the 2nd major cell type (Fig.1C), organoids containing only CC10+ve cells are rarely found in the culture. It appears that CC10+ve cells do not respond to KFH stimulation, and the culture conditions further enrich AT2 cells whilst reducing club cells.

Second, lineage tracing analysis has demonstrated that, in addition to AT2 cells, BASCs at the BADJ are also involved in lung regeneration (Liu et al. 2019; Salwig et al. 2019). Nevertheless, we observed very few BASCs in bronchioalveolar organoids. Our *in vitro* data support the notion that BASCs could restore both bronchial and alveolar epithelia after injury (Salwig et al. 2019), by giving rise to AT2 and club cells, and forming bronchioalveolar organoids (Fig.4I’,I”) (McQualter et al. 2010; Ng-Blichfeldt et al. 2018). It is plausible that, in culture, BASCs undergo asymmetric cell division to generate either proSPC+ve or CC10+ve positive daughter cells. Therefore, it would be of particular interest to identify the molecular mechanism that determines the fate of BASCs. We envisage that the organotypic culture method employed here could serve this purpose.

## EXPERIMENTAL PROCEDURES

### Mice

Colony maintenance of C57BL/6JRj mice (Janvier Labs) and experiments on female animals were performed in accordance with Directive 2010/63/EU and the regulations of the relevant local authorities (TLV, Thüringen, Germany; Ministry for Superior Education and Research, France), under the oversight of the FLI and PHENOMIN-ICS Animal Welfare Committees, including approval of the local Ethics Committee where appropriate.

The mice were provided with standard laboratory chow, tap water *ad libitum*, and they were kept at 22°C constant temperature and 12hr-light/12hr-dark light cycle. For generation of WT^Ad-SPC-Cre^, seven-week-old females were treated with Cyclosporine A (Neoral Novartis Pharmaceuticals) orally in the drinking water at a dose of 100mg/ml, one week prior to adenovirus administration and 2 weeks following adenovirus infection. Mice were intranasally administered with 20μl of purified Ad5-mSPC-Cre virus (2.10^8^ pfu). High titer adenoviruses (Ad5-mSPC-Cre) were either purchased (University of Iowa Gene Transfer Vector Core) or kindly provided by Anton Berns (NKI) (Ferone et al. 2016). Mice were sacrificed at 20 or 34 weeks of age for lung collection and AT2 cell isolation (Table S2).

### Antibodies used in cell isolation

Biotin-conjugated anti-CD45 (eBioscience, Cat# 13-0451, dilution 1/100); biotin-conjugated anti-EpCAM (eBioscience, clone G8.8, Cat# 13-5791-82, dilution 1/100); biotin-conjugated anti-T1α (Novus Biologicals, Cat# NB600-1015B, dilution 1/100).

### Organotypic culture of pulmonary epithelial cells

The cell pellet was re-suspended in 100% Matrigel to concentration of 6×10^6 cells/ml. The cell suspension was aliquoted (at 50μl/well) into an 8-well chamber slide (containing a 10mm x 7mm sterile plastic coverslip in each well) and incubated at 37°C in 5% CO_2_ for 30min, after which the solidified cell-Matrigel suspension was overlaid with AECC medium (400μl/well). The culture medium (which was renewed every second day) was supplemented with 10μM ROCK inhibitor (Y27632) during the first 2 or 4 days of culture (Fig.S1B), to aid cell attachment.

Organoids were allowed to form during 1 or 2 weeks in culture (Fig.S1B). On the 8^th^/14^th^ day, the medium was replaced first with PBS and then with 4% buffered paraformaldehyde (Kroll et al. 2016). The Matrigel-embedded organoids were fixed within the chamber slide for 48hr at RT; they were subsequently transferred to a histology cassette and immersed in 4% paraformaldehyde to continue fixation for another 48hr. Following dehydration in serial concentrations of ethanol, the organoids were embedded in paraffin and processed for immunohistochemistry.

### Transcriptome analysis

For transcriptome analysis, 5×10^5 EpCAM+ve cells were lysed in 350μL RTL Buffer (Qiagen) containing 143mM β-Mercaptoethanol (aided by cell searing via passage, 5 times, of the suspension through a 27G needle), snap-frozen in liquid nitrogen and stored at −80°C.

Total RNA extraction was performed with the AllPrep DNA/RNA kit (Qiagen) according to the manufacturer’s instructions. The integrity of extracted RNA was analyzed using the Bioanalyzer 2100 (Agilent). RNA libraries were constructed using the Illumina TruSeq v3.0 protocol, including RiboZero depletion, and sequenced on an Illumina HiSeq2500.

The same computational protocol was applied on alveolar cell and lung data. RNA reads were aligned to mouse reference mm10 using BWA (version0.7.5a). Gene counts based on Ensembl v87 gene models were calculated using custom scripts. Gene expression levels were quantified in reads per kilobase of exon per million mapped reads (RPKM). Differential gene expression was calculated using the Bioconductor package EdgeR, employing R-3.3.2.

### Proteome analysis

For proteome analysis, 1×10^7 EpCAM+ve cells were prepared for mass spectrometric analysis by lysis with 8 M urea and 0.1 M ammonium bicarbonate and tryptic digestion overnight into peptides. For phosphoproteome analysis samples were further enriched for phosphopeptides using TiO_2_ beads.

#### Mass Spectrometric Acquisition

For data dependent acquisition (DDA), 2 μg of peptides were injected to an in-house packed C18 column (Dr. Maisch ReproSil Pur, 1.9 μm particle size, 120 Å pore size; 75 μm inner diameter, 50 cm length, New Objective) on a Thermo Scientific Easy nLC 1200 nano-liquid chromatography system connected to a Thermo Scientific Q Exactive HF mass spectrometer equipped with a standard nano-electrospray source. LC solvents were A: 1 % acetonitrile in water with 0.1 % FA; B: 15 % water in acetonitrile with 0.1 % FA. The nonlinear LC gradient was 1 - 52 % solvent B in 60 minutes followed by 52 – 90 % B in 10 seconds, 90 % B for 10 minutes, 90 % - 1 % B in 10 seconds and 1 % B for 5 minutes. A modified TOP15 method from Kelstrup et al. (2012) was used. Full MS covered the m/z range of 350-1650 with a resolution of 60’000 (AGC target value was 3e6) and was followed by 15 data dependent MS2 scans with a resolution of 15’000 (AGC target value was 2e5). MS2 acquisition precursor isolation width was 1.6 m/z while normalized collision energy was centered at 27 (10% stepped collision energy) and the default charge state was 2+. Data-independent acquisition (DIA) was performed at the same LC-MS setup. A DIA method with one full range survey scan and 14 DIA windows was used, adopted from Bruderer et al. 2015. For measurements of phosphopeptide enriched samples 23 DIA windows method was used.

#### Mass Spectrometric Data Analysis

For generation of spectral libraries for DIA searches the DDA and DIA data were analyzed with the Pulsar search engine using SpectroMine™ v1 (Biognosys) applying the default settings. Search criteria included carbamidomethylation of cysteine as a fixed modification, oxidation of methionine and acetyl (protein N-terminus) as variable modifications and additionally phosphorylation of S, T, Y as variable modification for phosphopeptide enriched samples only. The DDA and DIA files were searched against the Mus Musculus Ensembl v87 database and the Biognosys’ iRT peptide sequences (Escher et al. 2012). The identifications were filtered to obtain a false discovery rate (FDR) of 1 % on peptide and protein level.

The DIA data was analyzed using Spectronaut™ v13 software (Biognosys) applying default settings. The FDR on peptide and protein level was set to 1 %. Local normalization was performed on the peptide quantities to compensate for loading differences and spray bias (Callister et al. 2006). Top3 peptides were used for calculation of protein quantities in total proteome data. DIA-specific PTM site localization score in Spectronaut was applied for the analysis of phosphopeptide enriched samples and phosphopeptides with score > 0.75 were reported.

#### Mapping mouse and human proteins

UniProt gene identifiers of phosphorylated proteins reported to be significantly changed after SARS-CoV-2 infection in C. sabaeus vero E6 cells (Bouhaddou et al. 2020) were mapped to Ensembl mouse gene identifiers to compare with AT2 cell data. Sequence positions of relevant phosphorylation sites were converted from human to mouse using PhosphoSitePlus® (Hornbeck et al. 2015).

### Gene Set Enrichment Analysis (GSEA)

GSEA was performed using gsea2-2.2.2.jar (Subramanian et al. 2005). Expression data was first selected for the subset of genes for which the protein was measured by MS and for genes with a minimal expression of 0.5 RPKM. Orthology mapping between mouse and human was performed using Ensembl v87 orthology tables. GSEA was performed on a pre-ordered list based on a ranking of isolated cells *versus* lung tissue with the distance D and fold-change FC by the metric: Ranking=sqrt(log10(D)^2 + log2(FC)^2) * sign(log2(FC)).

Further GSEA was performed using the gene sets h.all.v5.2, c2.cp.biocarta.v5.2, c2.cp.kegg.v5.2, and c2.cp.reactome.v5.2.

### Distribution of Materials & Data

Transcriptome and proteome files are provided as Supplementary Tables S2 & S3. Sequencing data have been deposited at the European Nucleotide Archive (ENA) under accession number PRJEB35893.

## ACKNOWLEDGEMENTS

The support of the Animal House, Histology, Imaging and Functional Genomics Core Facilities of the FLI as well as of the Animal House of PHENOMIN-ICS is gratefully acknowledged; in particular the expert assistance at FLI of Jenny Buchelt, Hellen Ahrens & Sabine Landmann and at PHENOMIN-ICS of Emilie Peter, Solene Schmitt & Peggy Mellul.

This study was funded by the European Union Horizon 2020 research and innovation program under grant agreement No. 686282, as part of the CanPathPro project. The FLI is a member of the Leibniz Association and is financially supported by the Federal Government of Germany and the State of Thuringia.

## SUPPLEMENTAL INFORMATION

### SUPPL. EXPERIMENTAL PROCEDURES

#### Reagents & materials used in cell isolation

50U/ml Dispase II in PBS (Sigma-Aldrich, Cat# 4942078001), aliquoted, stored at −20°C; DNase I (Roche, cat # 11284932001) stock concentration: 10mg/ml in Dulbecco’s-PBS; 1% low melting agarose (Sigma-Aldrich, Cat# A9419) prepared just before use; 40μm and 70μm cell strainer (BD Falcon); labeling buffer: 0.5% BSA, 2mM EDTA in PBS, filtered through 0.22μm filter, stored at 4°C; 10x red blood cell lysis buffer: 1.5M NH_4_Cl, 120mM NaHCO_3_, 1mM EDTA, filtered, stored at 4°C (diluted to 1x with sterilized ddH_2_O just before use).

Streptavidin-conjugated microbeads (Cat# 130-048-102); columns (Cat# 130-042-201); MiniMACS Separator (Cat# 130-042-102); 20μm pre-separation filter (Cat# 130-101-812); MACS MultiStand - all from Miltenyi Biotec.

For antibodies’ information, see Experimental Procedures in main section.

#### Culture media & supplements

MTEC basic medium (Lam et al. 2011): DMEM/F12 (ThermoFisher, Cat# 11330032), supplemented with 15mM HEPES, 2.5mM L-glutamine, 3.6mM sodium bicarbonate, 100U/ml penicillin, 100μg/ml streptomycin, 0.25μg/ml amphotericin B.

MTEC proliferation medium (Lam et al. 2011): MTEC basic medium supplemented with 5% FBS, 25ng/ml epidermal growth factor, 30μg/ml bovine pituitary extract, 0.1μg/ml cholera toxin, 1xITS-X, 0.01μM retinoic acid (filtered through a 0.22μm filter and added shortly before medium use).

Alveolar epithelial cell culture (AECC) medium was prepared using MTEC proliferation medium supplemented with one of the following component combinations:

i. Wnt5a (100ng/ml)
ii. Wnt5a (100ng/ml), CHIR99012 (50nM)
iii. KGF (50ng/ml), FGF10 (50ng/ml), HGF (40ng/ml)
iv. KGF (25ng/ml), FGF10 (25ng/ml), HGF (20ng/ml).

Y27632 (10uM) was included in the AECC medium during the first 2 or 4 days of culture (Fig.S1).

Culture supplements were obtained from the following providers:

**Table S4.**

| *Reagent* | *Supplier* | *Catalogue Number* |
| --- | --- | --- |
| Bovine pituitary extract | Corning | Cat# 354123 |
| CHIR-99021 | Selleckchem | Cat# S2924 |
| Cholera toxin | Sigma-Aldrich | Cat# C8052 |
| EGF | Sigma-Aldrich | Cat# E9644 |
| FGF10 | R&D systems | Cat# 6224-FG-025 |
| Growth-factor-reduced Matrigel | Corning | Cat# 354230 |
| HGF | Millipore | Cat# GF116 |
| Insulin-Transferrin-Selenium-X (ITS-X) 100x | ThermoFisher | Cat# 51500-056 |
| KGF | R&D systems | Cat# 5028-KG-025 |
| Retinoic acid | Sigma-Aldrich | Cat# R2625 |
| Wnt5a | R&D systems | Cat# 645-WN-010 |
| Y27632 | Selleckchem | Cat# S1049 |

#### Isolation of AT2 cells by magnet-associated cell sorting

The indicated quantities of materials are for AT2 cell isolation from one mouse. The procedure duration is *ca*. 6 hours. Mice were euthanized by cervical dislocation, followed by *in situ* lung tissue digestion, lung resection, further tissue digestion and preparation of single cell suspension as previously described (Gereke et al. 2012; Sinha and Lowell, 2016a; Sinha and Lowell, 2016b). After red-blood cell lysis, the cell pellet was re-suspended in 500μl labelling buffer containing 5μl DNase I and the cells were processed for the two-step magnet-associated cell sorting as follows.

##### Negative selection (Fig.S1A)

Biotinylated anti-CD45 and anti-T1α antibodies were added (5μl of each) to the cell suspension, which was then incubated (4°C, 30min) with intermittent mixing, to facilitate antibody-antigen binding. Unbound antibodies were diluted by addition of 10ml DMEM/F12 to the suspension; the cells were pelleted (200 x g, 10min, 4°C) and re-suspended in 90μl labeling buffer. Streptavidin-conjugated microbeads (10μl) and DNase I (1μl) were added and the suspension was incubated (4°C, 15min) with intermittent mixing. The cells were washed as above, and then re-suspended in 500μl labeling buffer. The cell suspension was loaded onto the column (which had been assembled on the MACS stand with the magnetic separator, and rinsed with 500μl labeling buffer prior to cell loading) through the 20μm pre-separation filter, to remove any large cell clumps. Filter and column were rinsed with 500μl labeling buffer and the unlabeled cell fraction, passing through the column, was collected. Filter and column were washed with 500μl labeling buffer, three times; the effluents were collected, pooled with the unbound cell fraction, then the cells were pelleted (200 x g, 10min, 4°C) and processed for positive selection, as described below. The column-bound cell fraction (CD45+ve and T1a+ve cells) was eluted, via removal of the column from the magnetic separator, for analysis.

##### Positive selection (Fig.S1A)

The cell pellet was re-suspended in 100μl labeling buffer, biotinylated anti-EpCAM antibody (1μl) and DNase I (1μl) were added and the suspension was incubated (4°C, 30min) with intermittent mixing. Unbound antibodies were diluted by addition of 10ml DMEM/F12 to the suspension; the cells were pelleted (200 x g, 10min, 4°C) and re-suspended in 90μl labeling buffer. Streptavidin-conjugated microbeads (10μl) and DNase I (1μl) were added and the suspension was incubated (4°C, 15min) with intermittent mixing. The cells were washed as above, and then re-suspended in 500μl labeling buffer. A new column (assembled on the MACS stand with the magnetic separator) was rinsed with 500μl labeling buffer, and loaded with the cell suspension. The unlabeled cell flow-through was collected for analysis. The column was washed with 500μl labeling buffer, three times; removed from the magnetic separator, placed into a 15ml tube and 1ml labeling buffer was pipetted into the column. The labeled cells were flushed from the column, using the plunger provided, and collected into the tube. The cells in the suspension were counted and processed for further analysis, or pelleted (200 x g, 10min, 4°C) and processed for organotypic culture.

#### Immunofluorescence microscopy & HCA of pulmonary cells

*Ca*. 200,000 EpCAM+ve cells were spun-down (2,000rpm, 5min, RT) onto poly-L-Lysine-coated slides (prepared according to Kroll et al. 2016) using a cytocentrifuge (Cytospin 4, Thermo Scientific). The cells were fixed by immersion of the slide in 3% paraformaldehyde (Kroll et al. 2016) for 10min, followed by permeabilization in −20°C methanol. Immunofluorescence labeling was performed according to Li et al. (2016) using the antibodies described under *Histochemistry and immunofluorescence microscopy of organoids & lung*.

For quantification of cell subtypes by HCA, tiled images of the entire cell monolayer, span-down on the slide, were acquired using a 10x objective in an Axiovert 200 microscope equipped with 12-bit grayscale cooled CCD AxioCamMRm camera (Zeiss). The tiled images were automatically stitched to a single image per monolayer imaged, during image acquisition (Zeiss ZEN software). Four representative fields-of-interest, cropped from each tiled image, were used for quantification of cell subtypes using Cell Profiler (proSPC, CC10, p63) (McQuin et al. 2018) or ImageJ (AQP5) (Schindelin et al.2012; Rueden et al. 2017).

Quantification of proSPC and CC10 labelled cells was performed on an average of 11000 cells per replicate (n=3) and cell fraction (EpCAM+ve/-ve); of p63 and CC10 labelled cells on an average of 5800 cells per replicate (n=3) and cell fraction (EpCAM+ve/-ve cell fraction); of AQP5 and proSPC labelled cells on average of 2600 cells per replicate (n=2) (EpCAM+ve fraction).

Maximum and average intensity per cell was measured in Cell Profiler; the cell types were first classified according to thresholds set based on the distribution of average intensities (Fig.1C). Following visual inspection by two operators (all cells per subtype group were scored up to a total of 90 cells/subtype; for populations comprising >90 cells, 90 randomly selected cells were scored), resulting correction estimates (0.6% to 5.5%) were applied to obtain the final classification (Table S5). The data is described in the Results as mean of the percentage of each cell subpopulation ±SD.

#### Histochemistry and immunofluorescence microscopy of organoids & lung

De-paraffinized and rehydrated 4μm thick sections of formalin-fixed paraffin-embedded lung tissue or organoids were processed for histochemical staining or immunofluorescence labelling according to Li et al. (2016) except for labelling with anti-proSPC (antigen retrieval at 95°C).

The following antibodies were used, in the indicated dilutions:

**Table S6.**
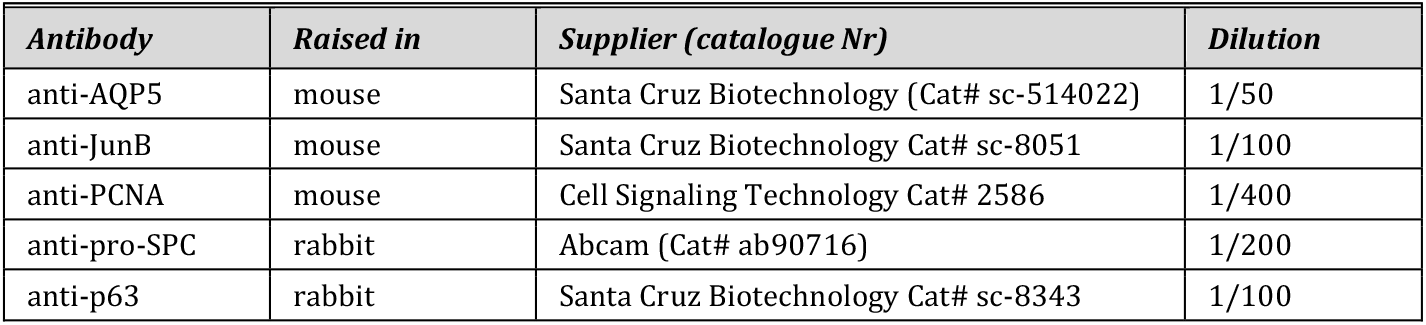

Microscopy of the histochemically-stained specimens employed a transmission light microscope (Olympus BX61VS) and images were acquired with a 20x objective using VS110 software (Olympus). Microscopy of the fluorescently-labeled specimens was performed according to Li et al. 2016.

#### Sample preparation for proteome analysis

Cell pellets were lysed with 8M urea and 0.1M ammonium bicarbonate. Protein concentration was determined by the mBCA assay (Pierce). 50μg protein per sample was prepared for mass spectrometry. The samples were reduced with 5mM tris(2-carboxyethyl)phosphine for 1h at 37 °C. Subsequently, the lysate was alkylated with 25mM iodoacetamide for 30min at 21°C. The lysate was diluted to 2M urea and digested overnight with sequencing grade modified trypsin (Promega) at 1:50 protease:protein ratio. The digested samples were desalted with C18 MicroSpin columns (SEM SS18V, Nest Group Inc.). Peptides were dried down using a SpeedVac system. The dried peptides were dissolved in LC buffer and iRT-peptide mix (Biognosys) was added for retention time calibration. Peptide concentrations were measured at 280nm with SPECTROstar® Nano spectrophotometer (BMG Labtech).

Pooled peptide samples were additionally fractionated for DDA analysis and spectral library generation using high pH reverse-phase fractionation. 60 μg peptides per pool were used. Ammonium hydroxide was added to a pH value > 10. The fractionation was performed using C18 MicroSpin columns (The Nest Group). Fractions were obtained by elution with increasing acetonitrile concentrations (5, 10, 15, 20, 25 and 50%). Fractions were dried down and resolved in Biognosys’ LC solvent. Fractions 5 and 50% were combined.

For phospho-proteome analysis, 1mg protein per sample were digested as above. Samples were desalted with C18 SepPak 100mg columns (Waters). Peptides were dried down to complete dryness using a SpeedVac system. Dried down peptides from cell lysates were redissolved in 50% lactic acid/50% acetonitrile/0.1% TFA solution and phosphopeptides were enriched using 10mg TiO2 beads (10μm, 300Å, Sachtopore, Sachtleben Chemie). Phosphopeptides were cleaned with C18 MicroSpin columns (The Nest Group). Dried down samples were redissolved with LC buffer and iRT-peptide mix.

## SUPPL. FIGURES & TABLES

**Figure S1:**
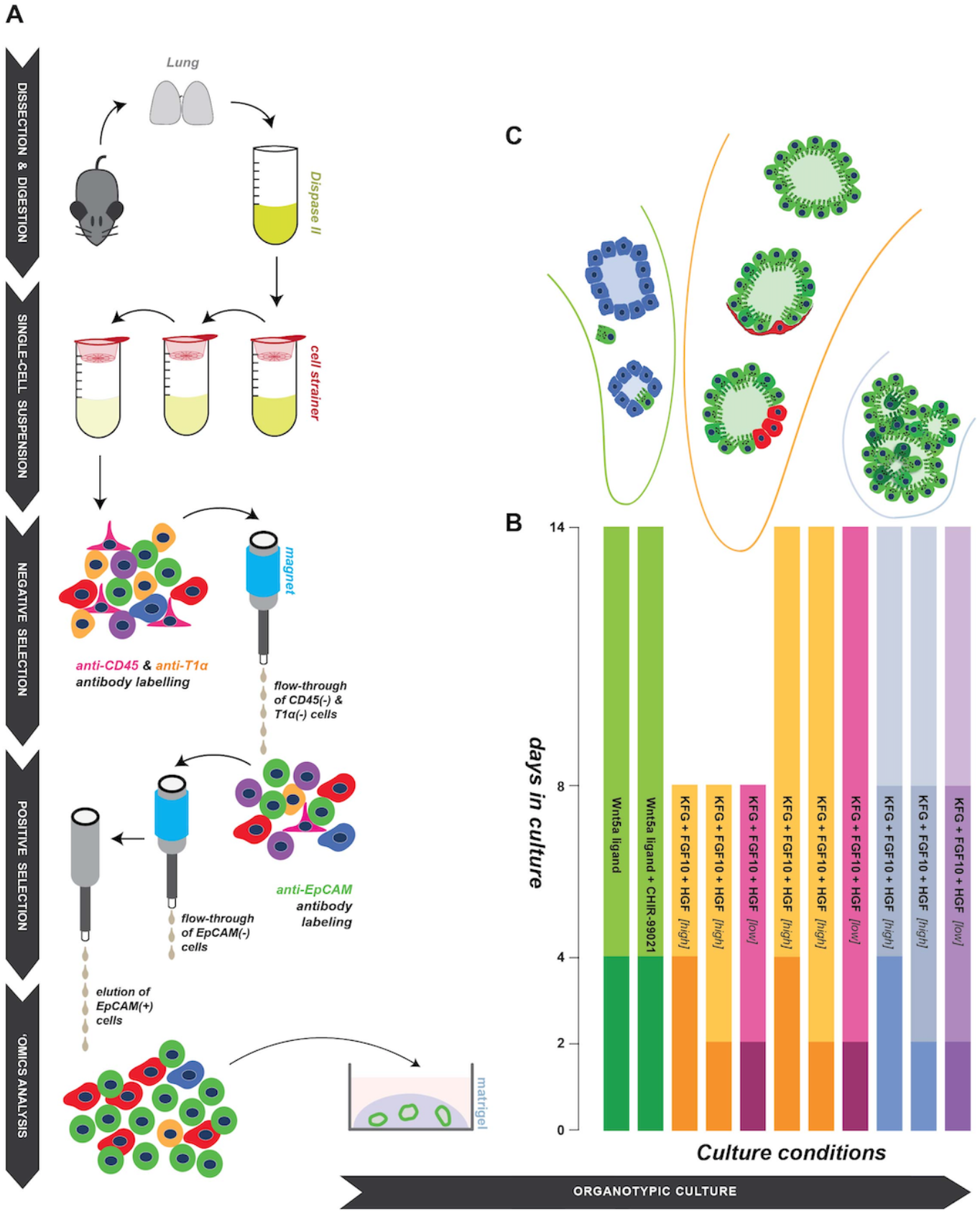
Schematic overview of AT2 cell isolation methodology, organotypic culture conditions and outcomes. (**A**) Dispase II digestion was initiated *in situ* and continued post lung dissection, according to Gereke et al. (2012), Sinha *et al.* (2016b). For enrichment of AT2 cells, by negative and positive selection and MACS-based separation, antibodies against surface proteins were utilized. Isolated EpCAM(+) cells were either ‘omics analyzed, or cultured to enable *in vitro* differentiation and organoid formation. (**B**) Schematic representation of the different culture supplement combinations tested for their ability to support AT2 organoid formation. The culture duration is indicated in the vertical axis. Each bar represents a culture condition, the darkest area of the bar indicates the period of Y-27632 supplementation (2-4 days). The lightest blue and purple bar areas indicate removal of KFH from the culture media (8-14 days). (See also *Experimental Procedures*). (**C**) Tracheospheres, bronchioalveolar organoids and alveolospheres were formed under the indicated culture conditions.

**Figure S2:**
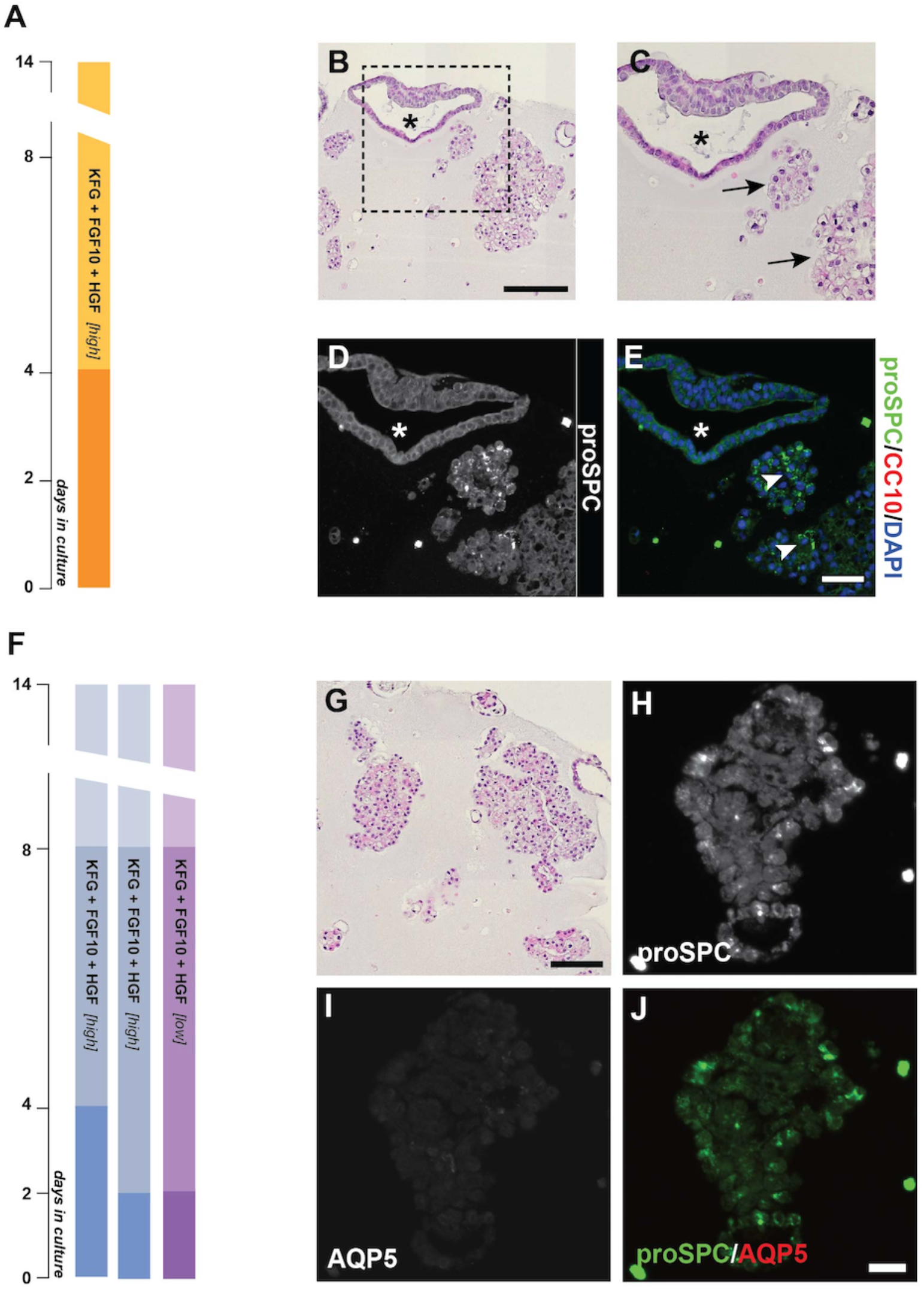
*In vitro* differentiation of AT2 cells. (**A, F**) Schematic representation of culture conditions (see also Fig.S1). The lightest bar areas, in **F**, indicate removal of KFH from the culture media (8-14 days of culture). H&E stained (**B, C**) and immunofluorescence labeled (**D, E**) semi-serial sections of 14-day organoids cultured as indicated in **A.** The asterisk marks a tracheosphere (**B-E**), arrows point to alveolar organoids (**C**) expressing proSPC (**E**, arrowheads). H&E stained (**G**) and immunofluorescence labeled (**H**-**J**) organoids cultured as indicated in **F** (high KFH). Large cell clumps of partly proSPC+ve cells are formed. Scale bars: 50μm (**E, H**), 100μm (**B**, **G**).

---

**Table S1:**
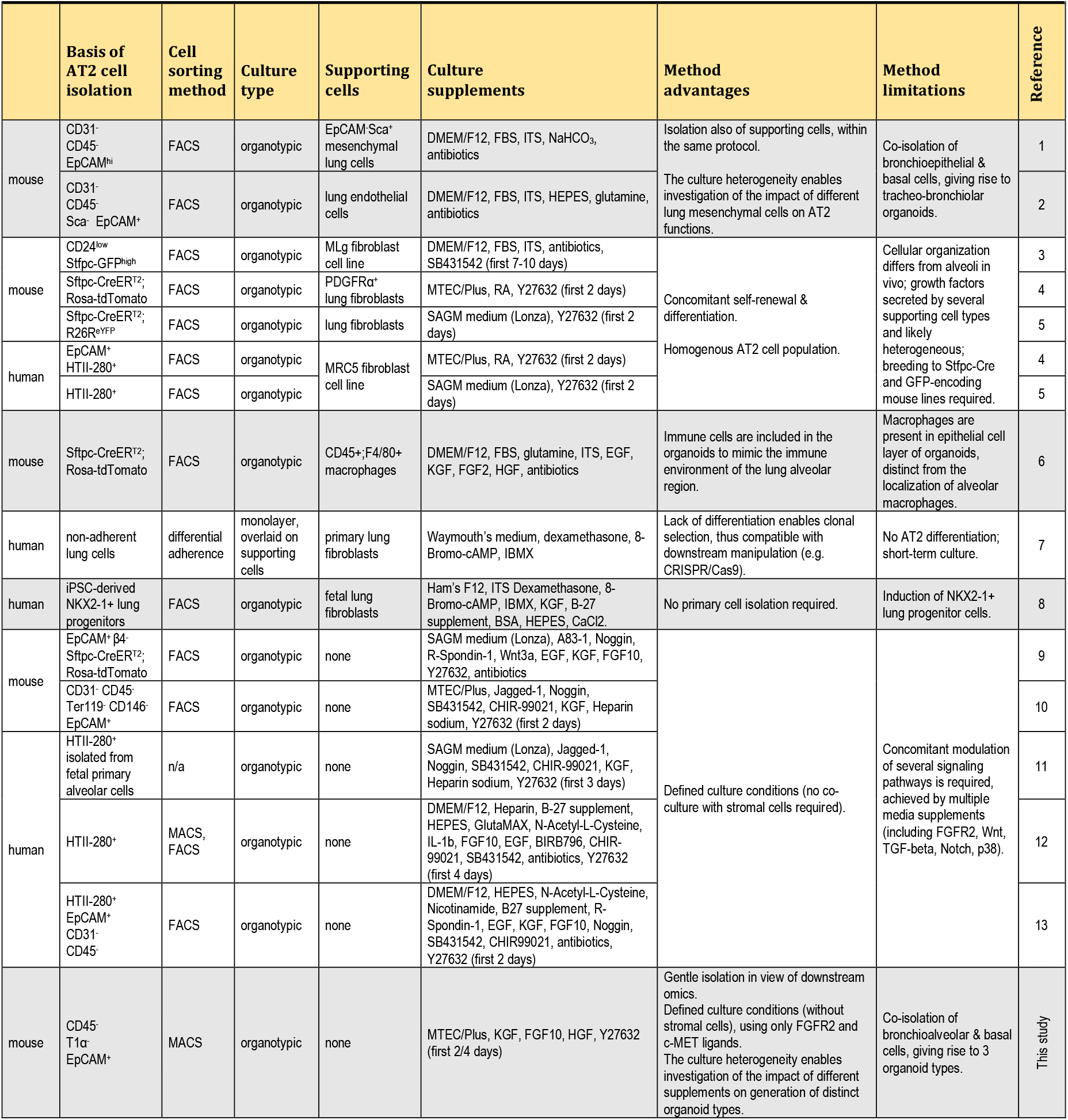
Overview of published methods used for the organotypic culture of AT2 cells. For the formulation of MTEC media, see Suppl. Experimental Procedures. References: (1) McQualter et al. 2010, (2) Lee et al. 2014, (3) Chen et al. 2012, (4) Barkauskas et al. 2013, (5) Zacharias et al. 2018, (6) Lechner et al. 2017, (7) Sucre et al. 2018, (8) Yamamoto et al. 2017, (9) Weiner et al. 2019, (10) Shiraishi et al. 2019a, (11) Shiraishi et al. 2019b, (12) Katsura et al. 2020, (13) Youk et al. 2020.

**Table S2:**

**Gene expression analysis (transcriptome and proteome) for lung tissue and EpCAM+ve pulmonary cell fraction.**

Gene expression data normalized as RPKM (columns H-P) and absolute label-free protein abundance (columns Q-Y) are shown.

**Table S3:**

**Expression and protein phosphorylation of selected components from the EpCAM+ve pulmonary cell fraction.**

**Sheet 1:** Gene expression and protein phosphorylation of (a) highly expressed AT2 cell-specific markers, (b) AP1 complex components which are enriched in the EpCAM+ve cell fraction, (c) putative AT2 cell surface markers. **Sheet 2:** Expression and protein phosphorylation of 40 SARS-CoV-2 interacting proteins (Bouhaddou et al. 2020), in the EpCAM+ve cell fraction. All detected phospho-sites are shown, with newly identified sites indicated in red and previously reported ones in black font.

**Table S5.**
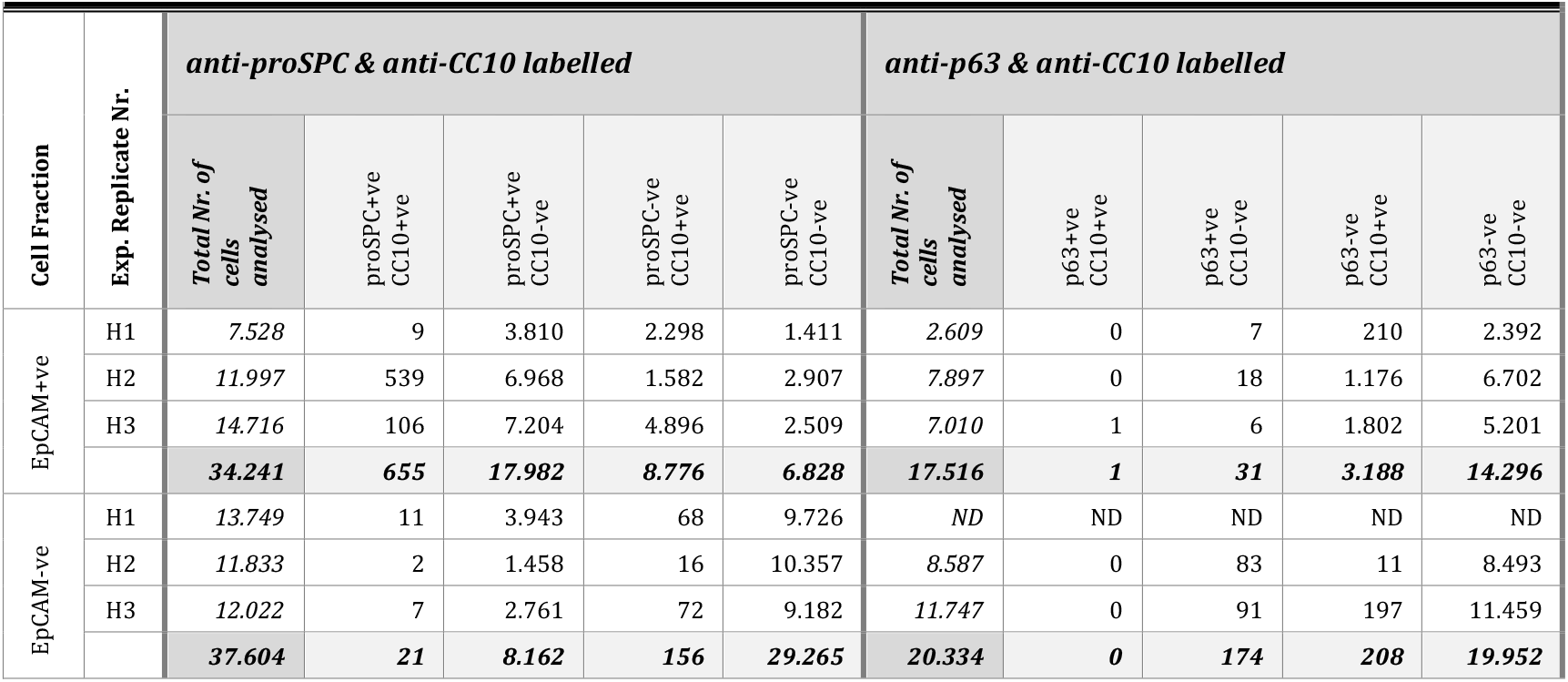

## Notes

### Competing Interest Statement

The authors have declared no competing interest.

